# Cell-specific α-tubulin TBA-6 and pan-ciliary IFT cargo RAB-28 generate a non-canonical transition zone

**DOI:** 10.1101/2023.11.16.567340

**Authors:** Jyothi Shilpa Akella, Malan S. Silva, Ken C. Q. Nguyen, David H. Hall, Maureen M. Barr

## Abstract

The transition zone (TZ) regulates cilia composition and function. Canonical TZs with 9 doublet microtubules (MTs) are common but non-canonical TZs that vary from 9 MT symmetry also occur and arise through unknown mechanisms. Cilia on the quadrant inner labial type 2 (IL2Q) neurons of *C. elegans* have a specialized non-canonical TZ with fewer than 9 doublet MTs. We previously showed that non-canonical TZs in IL2Q cilia arise via MT loss and reorganization of canonical TZs. Here, we identify structural events and mechanisms that generate non-canonical TZs. Cell-specific α-tubulin TBA-6 and pan-ciliary IFT cargo RAB-28 regulate IL2QTZ MT loss without affecting ciliary assembly. Our results reveal a role for the tubulin code in generating non-canonical TZs and contribute towards understanding ciliary functional specialization.

**Author summary:** Ciliary microtubules are exquisitely diverse in arrangements and composition. Studies on how ciliary ultrastructural diversity is generated are essential to our understanding of cilia function in diverse healthy and pathological contexts. Despite its clinical relevance, the ultrastructural diversity of the transition zone and its microtubules remains understudied. Here, we uncover mechanisms contributing to generating ultrastructural diversity in the transition zone and in cilia. A subset of sensory cilia in *C. elegans* contain a non-canonical transition zone with 7 and fewer doublet microtubules. We previously showed that this distinct transition zone is generated through microtubule loss in a canonical transition zone with 9 doublet microtubules, a process that occurs asynchronously during animal development. Here, we identify roles for the tubulin code and for an IFT cargo in generating a distinct transition zone. Sculpting of the distinct transition zone occurs in fully assembled cilia and transition zones and is independent of general ciliogenesis mechanisms. Our results demonstrate how specialized transition zones can be generated from canonical transition zones and provide insight into mechanisms of ciliary ultrastructural diversity and post-ciliogenesis restructuring. Such mechanisms hold the key to understanding ciliary function and to restoration of function in ciliopathies with ciliary ultrastructural defects.

## Introduction

Cilia are signaling organelles that are membrane-bound and contain a microtubule-based axoneme core. Intraflagellar transport (IFT) is a motor-based transport along ciliary microtubules (MTs) and builds most cilia (1). Cilia are ubiquitous and functionally diverse. Specializations in ciliary morphology, ultrastructure, and IFT facilitate context-specific functions of cilia. Defects in ciliary structure and ciliary function lead to disorders collectively termed as ciliopathies (2). Ciliopathies vary in their symptoms, which include blindness, kidney dysfunction, and altered cognitive abilities. Ciliary specialization mechanisms contribute to cell- and tissue-specific phenotypes of mutations in genes that encode cilia-localized products. Mechanisms of ciliary specialization include: (i) the tubulin code that consists of tubulin isotypes, tubulin post-translational modifying enzymes, motor proteins, and microtubule-associated proteins (ii) cell-specific transcription factors (iii) cell-specific regulation of IFT (3–11). Closing knowledge gaps in mechanisms of ciliary specialization is critical to our understanding of ciliary function and ciliopathies.

The transition zone (TZ) is the most proximal region of the axoneme and acts as a barrier to regulate ciliary composition and signaling functions (12–15). Mutations in genes encoding TZ-localized products are associated with human ciliopathies, highlighting the importance of the TZ to cilia function (16,17). The TZ is diverse in terms of composition, organization, and MT arrangements (18–20). A ring of 9 doublet MTs (dMTs) tightly connected to the ciliary membrane by characteristic Y-shaped linkers is the canonical form of TZ. However, non-canonical variations of the TZ that deviate from 9MT symmetry have been described in arthropods, apicomplexans, and nematodes (21–24). Although non-canonical TZs in mammalian cilia have not been reported yet, variations from 9MT symmetry are an evolutionarily conserved feature of cilia. Recent studies have identified non-canonical MT arrangements in other regions of the axoneme in epithelial, kidney, pancreatic islets, and glia and neurons in the brain (25–28). The functional impact and mechanisms of non-canonical ciliary MT arrangements, particularly in the TZ, remain largely unknown and are important to understand cell-and tissue-specific phenotypes in ciliopathies (29,30).

The nematode *C. elegans* presents an opportunity to identify mechanisms that regulate TZ specialization. Specifically, cilia on the inner labial type 2 (IL2) neurons have a specialized non-canonical TZ with 7 or fewer dMTs (31–33). We previously showed that the non-canonical TZ of IL2Q cilia arises from a canonical TZ (20). Generating a non-canonical TZ requires cell-specific reorganization involving MT loss and the equidistant rearrangement of the remaining MTs. IL2Q TZ reorganization occurs asynchronously during animal development and by adulthood, all IL2Q TZs are non-canonical. Mutants of genes encoding IFT-B components have defects in generating a non-canonical TZ in IL2Q cilia, indicating a role for IFT in IL2Q TZ specialization Here, we identify mechanisms that generate non-canonical TZs in IL2Q cilia.

Cell-specific and pan-ciliary molecules regulate ciliary ultrastructural and functional specialization. The IL2 neurons have a distinct tubulin code that includes a cell-specific α-tubulin isotype TBA-6, a cell-specific ciliary kinesin-3, and a cell-specific tubulin glutamylase: all components of the tubulin code (Figure 1A) (3,6,34,35). The IL2 neurons and four <u>ce</u>phalic <u>m</u>ale (CEM) neurons share these unique tubulin code components. In CEM cilia, TBA-6 sculpts non-canonical axonemes and influences extracellular vesicle (EV) shedding and signaling (34). Pan-ciliary molecules also regulate ciliary ultrastructural and functional specialization by performing cell-specific roles (7). RAB-28 is a pan-ciliary and membrane-associated IFT cargo that performs a specialized function in CEM neurons by regulating EV shedding (36). The roles of the tubulin code and RAB-28 in IL2Q TZ specialization remain unexplored.

**Figure 1.**
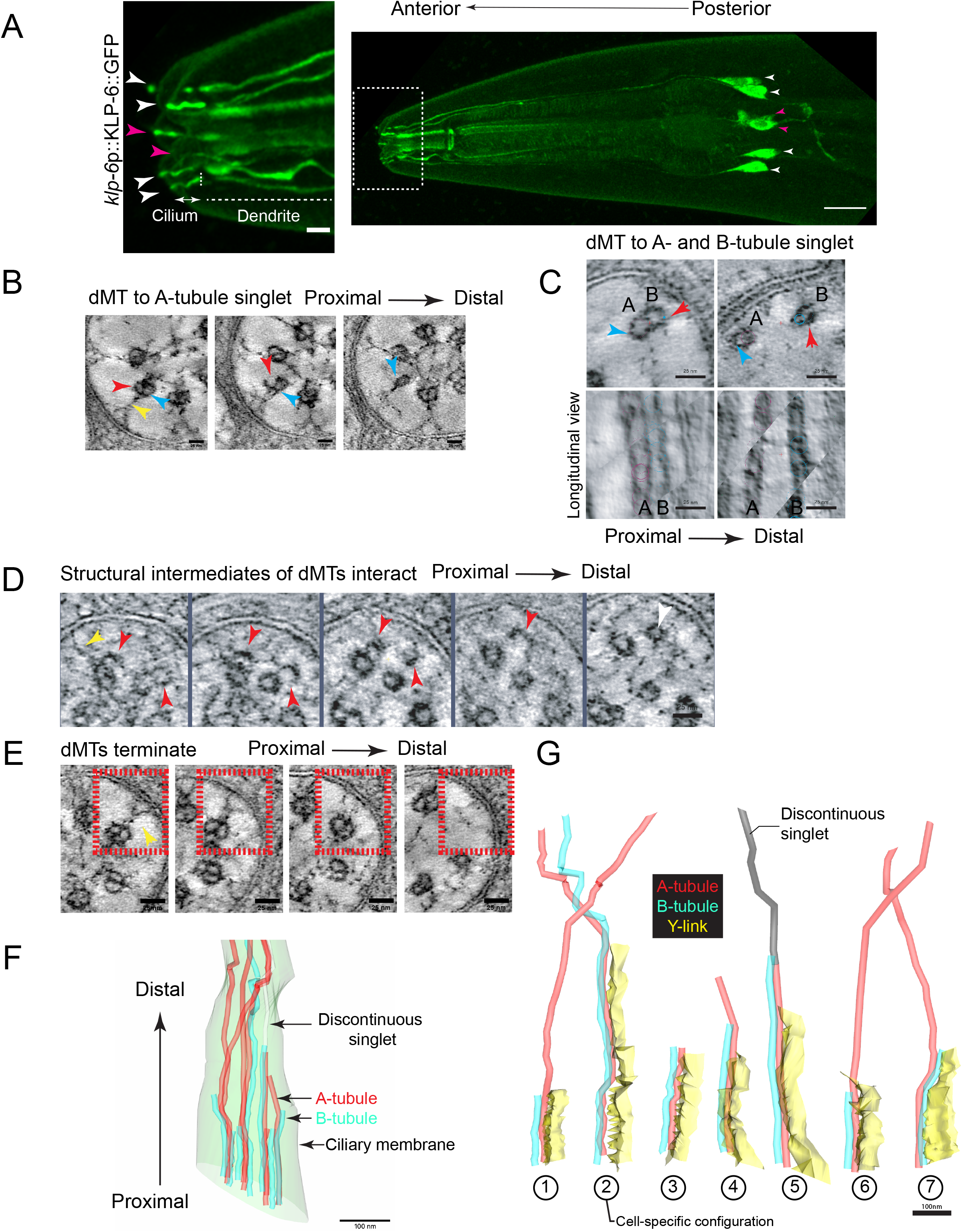
Ultrastructural characterization of IL2Q cilia and their MTs along their full length. (A) Confocal Z-projection of the IL2 neurons of *C. elegans* labeled using a *klp-6*p::KLP-6::GFP. White arrowheads mark the cell bodies of the quadrant IL2 (IL2Q) neurons that were studied in this paper; magenta arrowheads mark the cell bodies of the lateral IL2 (IL2L) neurons. The cilia are present on the tips of the neuronal dendrites. Both dendrites and cilia are marked and the demarcation between the dendrite and the cilium is represented by a vertical dotted line. Scale bar is 10µm. Area marked in inset is the nose of *C. elegans*. A closeup image of the area in the inset is shown on the left. White and magenta arrowheads in inset label IL2Q and IL2L cilia. All IL2 cilia are open to the environment through a pore in the cuticle. Scale bar is 2µm. (B) Slice-views of an ET following an IL2Q axonemal dMT in wild-type control *him-5* adults that adopts an A-tubule singlet configuration. Each image follows the same dMT in the proximal to distal direction. A-tubule, B-tubule, and Y-links are marked by cyan, red, and yellow arowheads, respectively. The B-tubule in the dMT disassembles leaving just the A-tubules distally. Scale bar is 25nm. (C) Slice-views of an ET following an IL2Q axonemal dMT in wild-type control *him-5* adults that displays a cell-specific configuration where both A- and B-tubules extend singlet MTs. The A-tubules (cyan arrowhead) and the B-tubules (red arrowheads) of the dMT split and persist as individual singlets more distally. Bottom panel shows a longitudinal view of the dMT before and after splaying. Scale bar is 25nm. (D) Slice-views of an ET following two IL2 axonemal dMT in wild-type control *him-5* adults that contribute to structural intermediates and discontinuous singlets. Shown here are partially disassembled B-tubules of adjacent MTs (red arrowheads) that interact with each other and appear as discontinuous singlet MTs (white arrowhead) more distally. Each image shows the same region of the cilium in the proximal to distal direction. Scale bar is 25nm. (E) Slice-views of an electron tomogram (ET) following an IL2 axonemal dMT in wild-type control *him-5* adults that gets terminated (MT enclosed in red box). Each image follows the same dMT in the proximal to distal direction. Yellow arrow points to Y-link in terminated dMT. Scale bar is 25nm. (F) 3D model of IL2Q cilia and their MTs. Model is based on serial tomogram of full length IL2Q cilia. MTs in IL2Q cilia display variability in lengths due to their disassembly and termination anywhere along the axoneme. A-tubules are marked in red, B-tubules in cyan, and discontinuous singlets are marked in white. The cilium is oriented longitudinally, so the proximal region is towards the bottom and the distal region is towards the top. Scale bar is 100nm. (G) 3D models of Y-linked dMTs in one IL2Q cilium. All MTs in this cilium are shown. Color scheme for A- and B-tubules is the same as in 1F, except for the discontinuous singlet, which is marked in grey here. Y-links are marked in yellow. B-tubule and Y-links terminate alongside each other in the A-tubule singlet configuration similar to amphid channel cilia. In the A-tubule and B-tubule singlet configuration similar to CEM cilia, Y-link termination occurs at the doublet splay point. Scale bar is 100nm.

Here we take advantage of the IL2Q cilia of *C. elegans* to study how non-canonical TZs are generated. Using transmission electron microscopy (TEM) and electron tomography (ET), we examine the ultrastructural basis of cell-specific reorganization of canonical TZs to generate a non-canonical TZ. We identify distinct features of IL2Q ciliary MTs, including the ability to assume multiple configurations and to undergo structural rearrangements along the axoneme. We observe cell-specific loss of TZ MTs and a coordination between MT and Y-link terminations in IL2Q cilia, identifying structural events associated with TZ reorganization. We discover a role for a cell-specific tubulin isotype TBA-6 in facilitating the MT loss of non-canonical TZs, the first identification of a role for tubulin isotypes in the ciliary TZ. Building on our previous observations on IFT associations of RAB-28 and its involvement with ciliary specialization mechanisms, we identify an IL2-specific role for RAB-28 in generating non-canonical TZs. RAB-28 regulates MT loss in IL2Q cilia without affecting the assembly of MTs with specialized features. Our findings establish the IL2Q cilia as a model to study mechanisms of TZ specialization and post-assembly structural reorganization in the cilium. Studies on ciliary structure may provide insight into cilia function and thus may be consequential to understanding ciliopathies.

## Results

### 1. Cell-specific MT loss and plasticity generate non-canonical TZs in IL2Q cilia

To understand how IL2Q cilia reorganize and generate a non-canonical TZ, we applied electron tomography (ET) to the entire length of the cilium. IL2Q ciliary MTs display unique features as well as similarities found in other cilia types in the *C. elegans* sensory nervous system. A subset of MTs in IL2Q cilia displayed an A-tubule singlet configuration where the B-tubules of the doublet MT terminated or disassembled and only A-tubules extended singlets distally (Figure 1B), resembling MT configurations of the amphid channel cilia (33). Some IL2Q ciliary MTs adopted a specialized configuration where A- and B-tubules of the dMT splayed and extended A-tubule and B-tubule individual singlets more distally (Figure 1C), similar to the <u>ce</u>phalic <u>m</u>ale-specific CEM cilia (34). These results suggest that MTs within a single IL2Q cilium are different from each other, contrasting with amphid and CEM cilia where MTs display a consistent architecture.

We identified specialized ultrastructural features unique to IL2Q cilia. We observed MT structural intermediates such as incomplete A- and B-tubules and noted interactions between structural intermediates (Figures 1B and 1D). Upon following the interactions between structural intermediates serially, we observed that structural intermediates appear as complete singlet MTs more distally. Some distal singlet MTs in IL2Q cilia do not trace back to a unique dMT or middle singlet originating in the TZ and are discontinuous, a deviation from other ciliary MTs (Figures 1D and 1F). Thus, IL2Q ciliary MTs are structurally plastic and not always extensions of MTs in the doublet of their origin. We also observed MT termination in IL2Q cilia (Figure 1E). Unlike MTs in amphid channel cilia that terminate at specific locations and mark transitions between distinct ciliary segments, MTs terminate anywhere along the IL2Q axoneme (Figure 1E and 1F). Thus, in addition to assuming different configurations, MTs are variable in lengths and origins. We determined if MTs displaying a specific configuration were targeted for loss and/or capable of structural plasticity. We found that MTs adopting A-tubule singlet configurations similar to amphid channel cilia (Figure 1B) and specialized A- and B-tubule singlet configurations similar to CEM cilia (Figure 1C) underwent disassembly and contributed to structural intermediates. Our data indicates that IL2Q-specific termination, disassembly, and structural plasticity of all MTs in IL2Q cilia results in deviations from canonical MT arrangements in TZ and in the rest of the axoneme.

After MT loss, Y-linked MTs remain equidistant from each other and retain symmetry in the TZ (20). Thus, despite being variable in configurations, lengths, and origins, MTs are symmetrically arranged within the non-canonical IL2Q TZ. To determine how this could be achieved, we examined MTs in the adult IL2Q TZ. We found that Y-link terminations largely correlated with B-tubule terminations in A-tubule singlet extensions and with the A-and B-tubule splay site in an A and B-tubule singlet extensions. In Figure 1G, MT# 1 is an example of an amphid-channel ciliary configuration with an A-singlet extension and MT#2 is an example of a CEM cilia configuration with an A-tubule and B-tubule singlet extensions. The correlation of Y-link terminations with MT terminations and transitions indicates a coordination between IL2Q ciliary MTs and Y-links to promote symmetry retention during reorganization of the TZ.

### 2. Cell-specific α-tubulin TBA-6 is required for non-canonical TZs in IL2Q cilia

To determine if the tubulin code generates non-canonical TZs in IL2Q cilia, we performed TEM on α-tubulin *tba-6* mutant adults (Figures 2A and 2B). TBA-6 is required for the A- and B-tubule singlet MT configuration in male-specific CEM cilia. In *tba-6(cxP4018)* mutant males, CEM cilia fail to splay dMTs into individual singlet MT extensions and instead display nine dMT extensions in the middle segment (34). In *tba-6* mutant IL2Q cilia, we consistently observed nine dMTs at the TZs (n=8/8), indicating that TBA-6 is essential for generating non-canonical TZs (Figures 2B and 2C). The presence of canonical IL2Q TZs in *tba-6* mutants indicates that another α-tubulin can substitute for TBA-6. This α-tubulin can also substitute for TBA-6 in IL2Q TZ elongation (Figure 2D). To determine if TBA-6 was required for IL2Q ciliary extension, we visualized IL2Q cilia in living wild-type and *tba-6* mutant animals using an IL2-specific ciliary reporter CIL-7::GFP (Figure 2E). In wild-type and *tba-6* mutants, IL2Q cilia elongated and protruded through the cuticle to the environment. We conclude that TBA-6 is required for IL2Q TZ MT loss, not TZ or ciliary extension.

**Figure 2.**
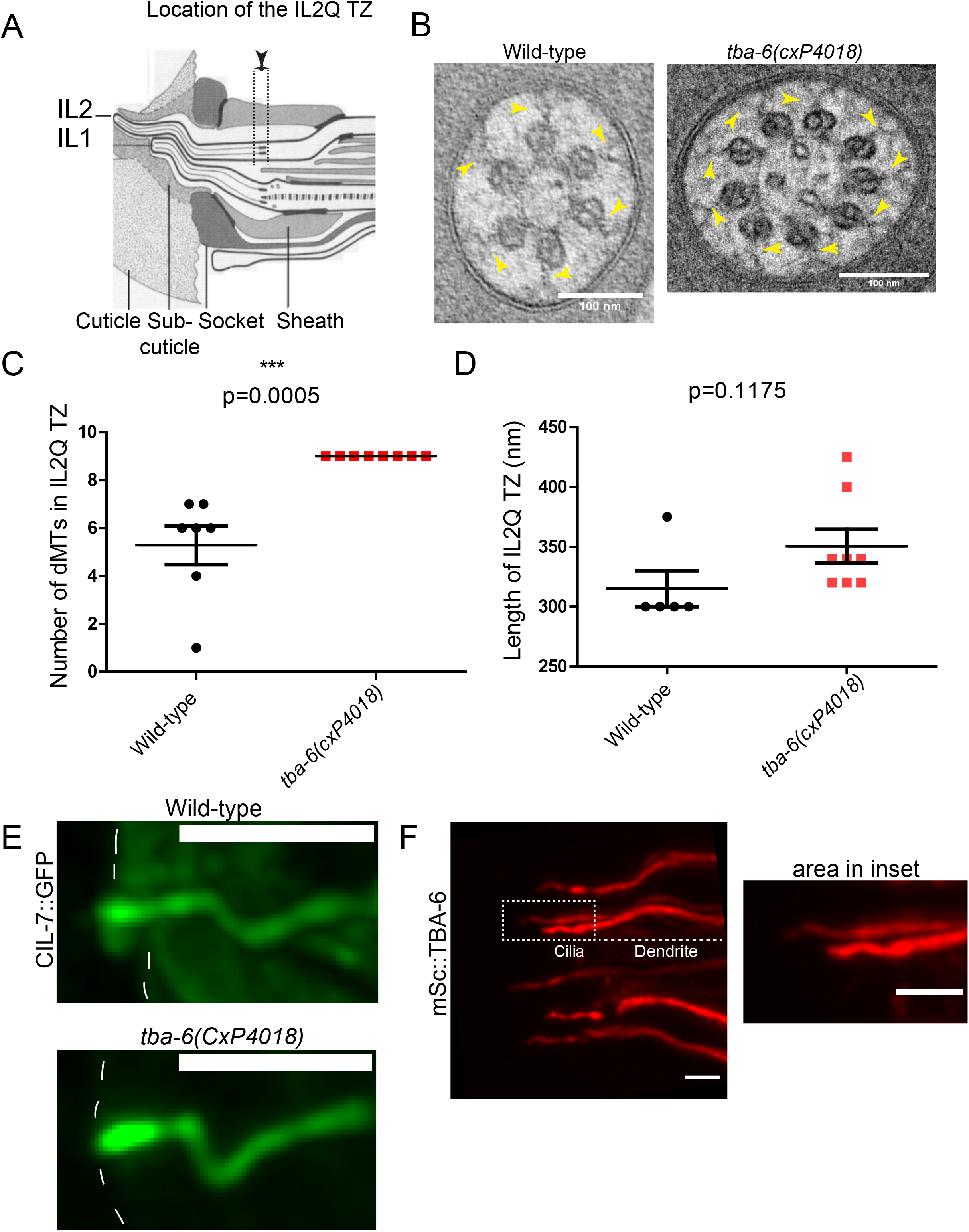
A cell-specific MT composition via TBA-6 regulates IL2Q TZ specialization. (A) Cartoon of the inner labial (IL) sensory organ of *C. elegans.* The IL sensory organ comprises the sensory endings of the IL1 and IL2 neurons that are engulfed by the IL sheath and socket glial cells, and the cuticle. The IL1 cilia remain embedded in the cuticle while the IL2 cilia poke out into the environment through a pore in the cuticle. Dotted lines mark the TZ region, the length of which was measured in D. Image reproduced from (31) (B) Cross-section TEM of the TZ of IL2Q cilia in wild-type control *him-5* and *tba-6(cxP4018)*; *him-5* adult males*. tba-6* mutants have a canonical TZ at adulthood. Y-links associated with MTs at the TZ are marked by yellow arrowheads. Scale bar is 100nm. (C) Scatter plot depicting the number of dMTs observed in the TZ of wild-type control *him-5* and *tba-6(cxP4018)*; *him-5* mutant adult male IL2Q cilia. *tba-6(cxP4018)*; *him-5* mutant IL2Q TZs contain more dMTs as determined by a Mann-Whitney test. p=0.0005; n=7 cilia and N=2 animals for wild-type control *him-5*; n=8 cilia and N=2 animals for *tba-6(cxP4018)*; *him-5*. (D) Scatter plot depicting the lengths of the TZs in IL2Q cilia of wild-type control *him-5* and *tba-6(cxP4018)*; *him-5* mutant adult male IL2Q cilia. Defects in IL2Q TZ specialization in *tba-6(cxP4018)*; *him-5* mutant adult males did not affect TZ extension as analyzed by an unpaired t-test with Welch’s correction. P=0.1175; n=5 cilia and N= 2 animals for wild-type control *him-5*; n=8 cilia and N= 2 animals for *tba-6(cxP4018)*; *him-5*. (E) Confocal Z-projections of wild-type control *him-5* and *tba-6(cxP4018)*; *him-5* mutant adult hermaphrodites expressing *cil-7p*::CIL-7::GFP. IL2Q cilia in TZ specialization defective *tba-6* mutants have no gross defects in ciliary extension and penetrate the cuticle to access the environment. The outline of the animal is marked by dotted lines. Scale bar is 2µm. (F) Confocal Z-projection of adult hermaphrodite *C. elegans* expressing endogenously CRISPR tagged mScarlet::TBA-6. Area marked in dotted white rectangle marks two IL2Q cilia that are shown in inset. TBA-6 localizes throughout the IL2Q cilium. Scale bar is 2μm.

To study endogenous TBA-6 protein localization in IL2Q cilia, we CRISPR edited the *tba-6* genomic locus to encode an N-terminal tagged reporter mScarlet::TBA-6. Endogenous TBA-6 was expressed in IL2 neurons, similar to published extrachromosomal and endogenous reporters (37,38). mSc::TBA-6 localized throughout IL2Q cilia (Figure 2F), suggesting that TBA-6 acts along the cilium and not at a specific ciliary region. mSc::TBA-6 localization is consistent with MT rearrangements and losses along the IL2Q cilium. Our data identifies a novel role for the tubulin code via cell-specific α-tubulin TBA-6 in generating non-canonical TZs.

### 3. Pan-ciliary IFT cargo RAB-28 generates non-canonical TZs in IL2Q cilia

The pan-ciliary GTPase RAB-28 plays a role in the ciliary specialization of the CEM neurons (36). RAB-28 is an IFT cargo in various cilia types in *C. elegans* (36,39). In IL2 neurons, mutations in some IFT-B genes result in a canonical 9dMT TZ (32). To determine if RAB-28 plays a role in generating non-canonical TZs in IL2Q cilia, we performed TEM on *rab-28(tm2636)* mutant adults. In *rab-28* IL2Q cilia, we observed 9dMTs at the TZs (Figures 3A and 3B). Unlike *tba-6(cxP4018)* mutants that completely lose the ability to generate a non-canonical TZ, the *rab-28(tm2636)* phenotypic defect was less severe. In *rab-28* adults, a canonical TZ was observed in 4 of 10 IL2Q cilia and a non-canonical TZ in 6 of 10 IL2Q cilia (Figure 3B). *rab-28* mutant cilia with non-canonical TZs displayed 7 or 8 MTs in IL2Q TZs (Figure 3B). Similar to *tba-6, rab-28* mutants were not defective in IL2Q TZ and ciliary extension (Figures 3C and 3D), indicating that mechanisms that generate non-canonical TZs are distinct from mechanisms regulating TZ and ciliary extension. Altogether, our results extend previous findings on cell-specific functions of RAB-28 and a relationship between IFT and IL2 TZ specialization.

**Figure 3.**
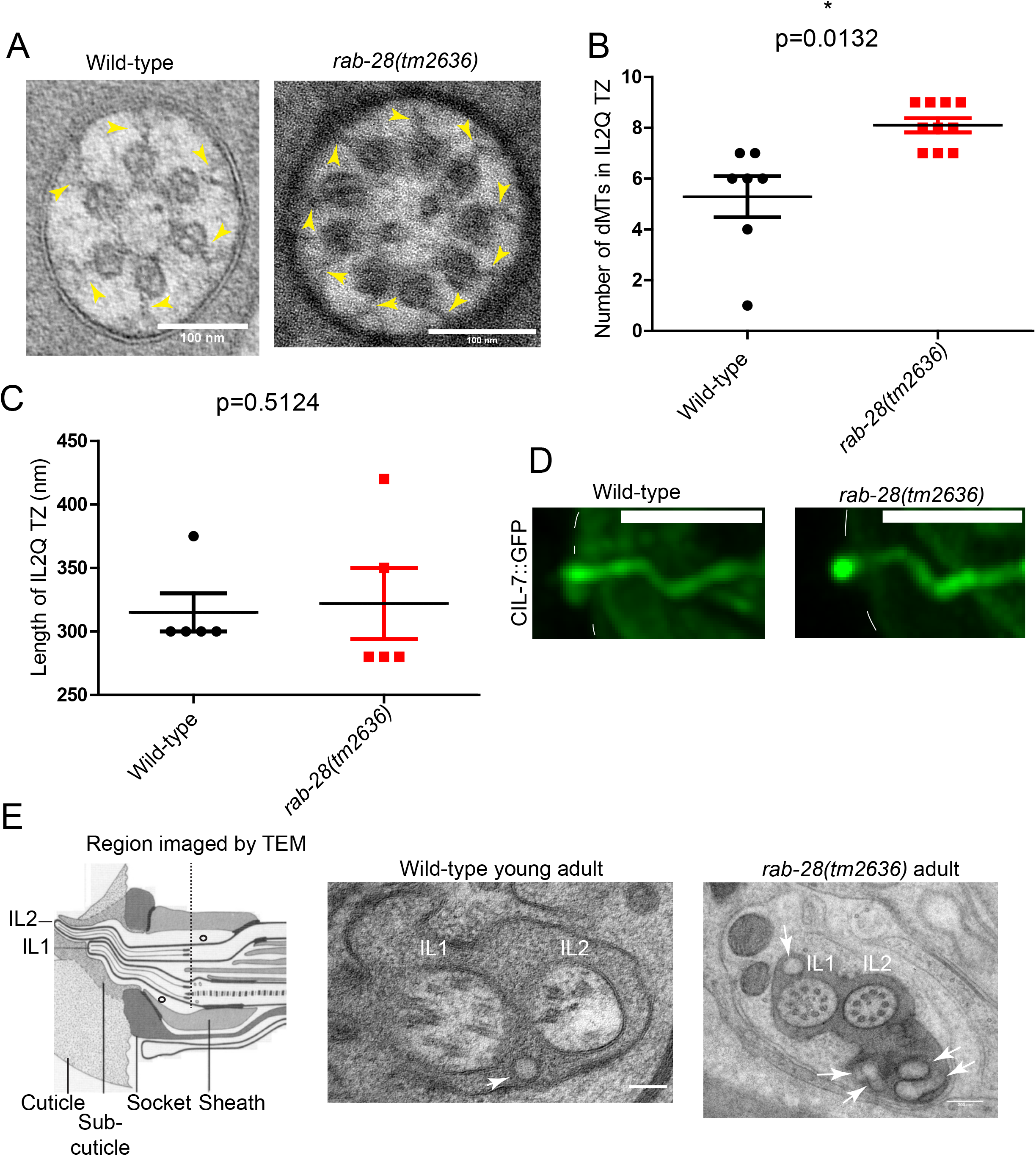
Pan-ciliary IFT cargo RAB-28 regulates IL2Q TZ specialization and membrane shedding in inner labial (IL) sensory organs. (A) Cross-section TEM of the IL2Q ciliary TZ of adult males of wild-type control *him-5* and *rab-28(tm2636)*; *him-5* mutants*. rab-28* mutants have more dMTs at the TZ. Y-links associated with MTs at the TZ are marked by yellow arrowheads. Scale bar is 100nm. (B) Scatter plot depicting the number of dMTs observed in the TZ of wild-type control *him-5* and *rab-28(tm2636); him-5* mutant adult male IL2Q cilia. *rab-28(tm2636)*; *him-5* mutant IL2Q TZs contain more dMTs as determined by a unpaired t-test with Welch’s correction. p=0.0132; n=7 cilia for wild-type *him-5*, and 10 cilia for *rab-28(tm2636)*; *him-5.* Data is from at least two animals for both genotypes. (C) Scatter plot depicting the lengths of the TZs in IL2Q cilia of wild-type control *him-5* and *rab-28(tm2636)*; *him-5* mutant adult male IL2Q cilia. Defects in IL2Q TZ specialization in *rab-28(tm2636)*; *him-5* mutant adult males did not affect TZ extension as analyzed by a Mann-Whitney test. P=0.5124; n=5 cilia and N= 2 animals for wild-type control *him-5* and n=5 cilia and N= 2 animals for *rab-28(tm2636)*; *him-5*. (D) Confocal Z-projections of wild-type control *him-5* and *rab-28 (tm2636)*; *him-5* mutant adult hermaphrodites expressing *cil-7p*::CIL-7::GFP. IL2Q cilia in TZ specialization defective *rab-28* mutants have no gross defects in extension and penetrate the cuticle to access the environment. The outline of the animal is marked by dotted lines. Scale bar is 2µm. (E) Left - Same cartoon as in Figure 2A. EVs (marked by circles) are shed within an extracellular lumenal space created by the surrounding glia. In inner labial sensory organs, EVs can be found throughout the lumen. Dotted line represents the region imaged in wild-type control and *rab-28(tm2636); him-5* mutants by TEM on the right. Right-Cross-section TEM of the TZ of a quadrant IL sensory organ of adult males of wild-type control *him-5* and *rab-28(tm2636)*; *him-5* mutant animals. White arrow in wild-type control panel is pointing towards a shed EV in the IL sensory organ. Scale bar is 100nm. In comparison to control *him-5* animals, *rab-28(tm2636)*; *him-5* mutant males accumulate an excess of membranous vesicles in the inner labial sensory organs (white arrows). Scale bar in *rab-28 (tm2636)*; *him-5* mutant panel is 200nm.

RAB-28 is also a negative regulator of ciliary EV shedding in CEM neurons (36). To determine if RAB-28 regulates EV shedding in the IL2Q neurons, we compared TEM images on the inner labial sensory organs of wild-type and *rab-28(tm2636)* adults. The IL2Q cilia are housed within inner labial sensory organs that contain a lumenal space where EVs are observed (Figure 3E) In the lumen of *rab-28* mutant inner labial organs, we observed excessive membranous accumulations not observed in wild-type adults (Figure 3E and Figure 3 figure supplement 1A and 1B). Thus, RAB-28 regulates EV shedding in both cephalic and inner labial sensory organs.

#### 1.4. RAB-28 regulates TZ MT loss in IL2Q cilia and not MT specialization

To understand how RAB-28 influences the formation of non-canonical TZs, we performed ET on the entire length of IL2Q cilia. In *rab-28 (tm2636)* IL2Q neurons, we observed cilia with dMTs that adopt an amphid-like A-tubule singlet configuration and dMTs that adopt a CEM-like A- and B-tubule singlet configuration (Figures 4A and 4B). We noted structural intermediates and interactions between structural intermediates, contributing to discontinuous singlets (Figure 4C). We also observed MT termination and MT disassembly in *rab-28* mutant IL2Q cilia, explaining how a subset of *rab-28* mutants have non-canonical TZs (Figure 4D). The retention of specialized IL2Q ciliary MT features in *rab-28 (tm2636)* mutants suggests that RAB-28 does not affect IL2Q ciliary MT specialization (different MT arrangements are quantified in Figure 4E). Compared to wild type, B-tubules in *rab-28 (tm2636)* IL2Q cilia extended farther past the Y-linked region, suggesting defects in correlating B-tubule and Y-link terminations (Figures 4F and 4G). The presence of normal length IL2Q TZs in *rab-28* mutants (Figure 3C) suggests that Y-link termination may be uncoupled from B-tubule termination and that RAB-28 regulates the latter. Despite defects in correlating Y-link and B-tubule terminations, *rab-28* mutants retained TZ symmetry in IL2Q cilia. Our data suggests that RAB-28 regulates MT loss in the TZ and B-tubule termination at and above the TZ without impacting IL2Q MT specialization.

**Figure 4.**
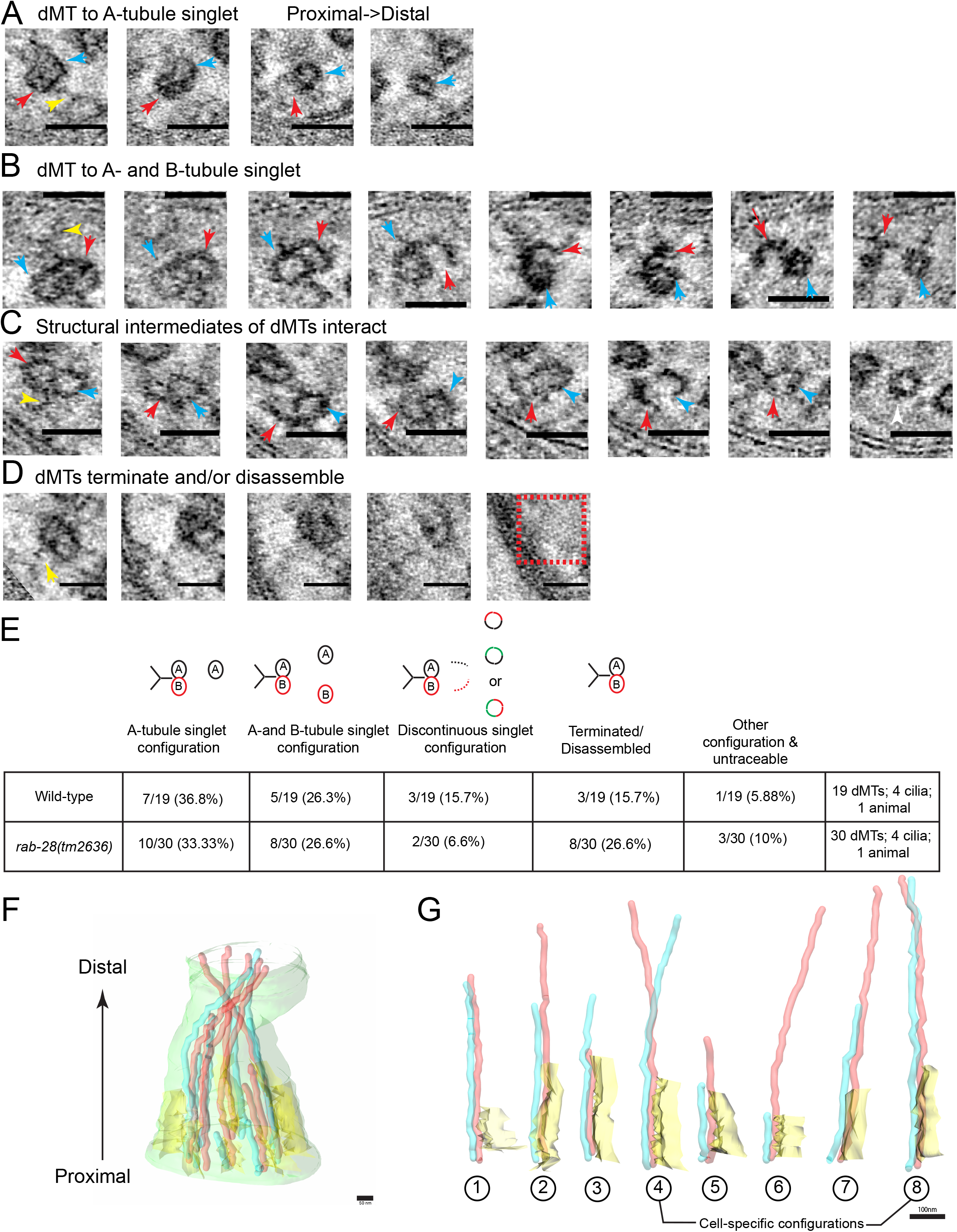
RAB-28 regulates IL2Q TZ specialization by controlling TZ MT loss and not MT specialization. (A) Slice-views of an electron tomogram (ET) following an IL2Q axonemal dMT in *rab-28(tm2636); him-5* adults that displays an amphid-type A-tubule singlet configuration. Each image follows the same dMT in the proximal to distal direction. The B-tubule (red arrowhead) of the dMT disassembles leaving only an A-tubule (cyan arrowhead) in the more distal regions, similar to amphid channel cilia. Yellow arrowhead marks the Y-link associated with the dMT. Scale bar is 25nm. (B) Slice-views of an electron tomogram (ET) following an IL2Q axonemal dMT in *rab-28(tm2636); him-5* adults that displays a CEM-type A- and B-tubule singlet configuration. Each image shows the same dMT in the proximal to distal direction. The A- and the B-tubules of the dMT (marked by cyan and red arrowheads, respectively) split and persist as individual singlets more distally. Yellow arrowhead marks the Y-link associated with the dMT. Scale bar is 25nm. (C) Slice-views of an ET following an IL2Q axonemal dMT in *rab-28* mutants that contribute to structural intermediates and discontinuous singlets. Each image shows the same dMT in the proximal to distal direction. The A- and the B-tubules of the dMT (marked with cyan and red arrowheads, respectively) partially disassemble and interact with each other and appear as a discontinuous singlet in anterior sections (white arrowhead). Yellow arrowhead marks the Y-link associated with the dMT. Scale bar is 25nm. (D) Slice-views of an electron tomogram (ET) following an IL2 axonemal dMT in *rab-28* mutants that gets completely disassembled. Region marked in red box indicates absence of the dMT more distally. Each image follows the same dMT in the proximal to distal direction. Yellow arrow points to Y-link in terminated dMT. Scale bar is 25nm. (E) Quantification of MT configurations observed in ET of IL2Q cilia of wild-type control *him-5* and *rab-28(tm2636)*; *him-5* adult males. *rab-28(tm2636)*; *him-5* IL2Q adopted all the MT configurations observed in control *him-5* cilia. Despite retaining more MTs at their TZ, *rab-28(tm2636)*; *him-5* MTs terminated and disassembled similarly to WT. Data is from n=19 dMTs from 4 cilia in 1 *him-5* control animal and n=30 dMTs from 4 cilia in 1 *rab-28(tm2636)*; *him-5* animal. (F) 3D model of IL2Q cilia and their MTs in *rab-28 (tm2636)* mutants. Model is based on serial tomogram of a full length IL2Q cilium. A-tubule, B-tubule, and Y-links are marked by cyan, red, and yellow arrowheads, respectively. The cilium is oriented longitudinally, so the proximal region is towards the bottom and the distal region is towards the top. Scale bar is 50nm. (G) 3D models of Y-linked dMTs in one IL2Q cilium in a *rab-28(tm2636)*; *him-5* mutant adult male. All MTs in this cilium are shown. Like wild-type, MTs in IL2Q cilia of *rab-28(tm2636)* are variable in length and configuration. B-tubules in MT#s 1,2,3, and 7 do not terminate closer to Y-link termination points. Color scheme is the same as in figure 1G.

We conclude that RAB-28 regulates the formation of non-canonical TZs by facilitating TZ MT loss and correlating Y-link and B-tubule terminations in axonemes with intact cell-specific MT arrangements.

## Discussion

### 1. Cell-specific MT composition is essential for generating non-canonical TZs in IL2Q cilia

Cell-specific α-tubulin isotype TBA-6 is necessary for IL2Q TZ specialization. MTs lacking TBA-6 fail to generate non-canonical TZs and default to a canonical TZ. The *C. elegans* genome encodes nine α-tubulins: six are expressed in ciliated neurons and three localize to cilia (38,40–42). Our data suggests that one of these six ciliated neuron-expressed α-tubulins may substitute for TBA-6 in ciliogenesis, but none can compensate for TBA-6 in TZ specialization. Tubulin isotypes generate MTs with distinct protofilament numbers and sculpt specialized axonemes (34,43–46). Our discovery of a cell-specific role for a tubulin isotype in IL2Q TZ specialization is consistent with established roles for tubulins in MT and ciliary specialization. Ciliary tubulin isotypes are essential in human development and health: mutations in a β-tubulin isotype cause human ciliopathies (47). Our results may provide an explanation for cell- and tissue-specific differences in the phenotypic severity of ciliopathies, including those caused due to mutations in tubulin encoding genes.

We hypothesize that isotype-specific MT dynamics play a role in generating non-canonical TZs in IL2Q cilia. MTs assembled using different tubulin isotypes differ in their growth and depolymerization rates, lifetimes, and catastrophe frequencies (11,34,43,44,46,48–51). Experiments in *C. elegans* involving β-tubulin isotype substitution and knockdowns to generate isotypically pure MTs revealed differences between the growth rates and lifetimes of MTs (50). Furthermore, the β-tubulin isotype incorporated within the polymer also influenced the effect α-tubulins had on MT growth rate (50). These *in vitro* and *in vivo* findings suggest that TBA-6-containing MTs may have different dynamics compared to MT assembled with other α-tubulin isotypes. TBA-6 has a longer C terminal tail (CTT) region that may interact with motors and microtubule-associated proteins (MAPs) (34). *In vitro* and cell culture-based studies show that different β-tubulin isotypes influence the action of depolymerizing MAPs on MTs (46,51). Thus, MTs assembled with TBA-6 versus other α-tubulins may differ in MAP-based modulation of dynamics and/or interactions with TBB-4, the ciliary β-tubulin expressed in IL2 neurons (34,37).

A striking feature of IL2Q ciliary MTs is the variability in length and ultrastructural configurations. Our finding that another α-tubulin(s) can substitute for TBA-6 in IL2 ciliogenesis indicates that IL2Q ciliary MTs are compositionally heterogenous and contain at least one other α-tubulin. Consistent with this idea, distinct tubulin isotypes may copolymerize into MTs and the tubulin requirements for MT polymerization *in vivo* display redundancy (34,43,44,46,48–51). We propose that individual MTs in IL2Q cilia differ in α-tubulin isotype content and undergo termination and disassembly at different rates, explaining asynchrony and variability between cilia (20) (Figures 1 and 4).

Future studies examining the relationship between TBA-6 and other components of the tubulin code such as other tubulin isotypes and MAPs in IL2Q neurons, will clarify how TBA-6 generates non-canonical TZs.

### 2. RAB-28-mediated IFT is important to generate non-canonical TZs

We show that IFT cargo RAB-28 is important to IL2Q TZ specialization, consistent with previous ultrastructural studies showing that IFT-B mutants fail to generate a non-canonical TZ (32). Several mutants associated with human ciliopathies (*che-13/*IFT57, *osm-5*/IFT88, *osm-6*/IFT52, and *osm-1*/IFT172) display 9-fold symmetric canonical IL2Q TZs as adults (32,53,54). Our findings are in agreement with the known roles for IFT in TZ gating and maintenance (7,55,56).

We hypothesize that IFT regulates IL2Q TZ specialization via a role in the transport of RAB-28, TBA-6, and yet-to-be identified TZ specialization regulators. Tubulin transport in cilia occurs both via IFT and by diffusion, supporting a role for IFT in TBA-6 transport (40,57,58). In *Chlamydomonas*, during ciliary disassembly, IFT plays a role in the transport of a microtubule depolymerizing protein kinesin-13 and a CDK-like kinase FLS2 that regulates kinesin-13 phosphorylation (59,60). IFT may similarly mediate the transport of critical MT regulators and/or the molecules that control the activity of MT regulators in IL2Q cilia. That *rab-28* mutants phenocopy the IL2Q TZ defects observed in IFT-B mutants suggests that *rab-28* and IFT-B genes act in the same genetic pathway. An interdependent relationship between RAB-28 and IFT may be essential for the maintenance of cell-specific functions of RAB-28 in IL2Q cilia. Determining the mechanism of RAB-28 action and identifying IFT cargoes will shed light on the role of IFT in IL2Q TZ specialization.

### 3. IL2Q cilia are a model to study ciliary plasticity

Cilia display plasticity - the capacity to change. One of the most consequential yet understudied forms of ciliary plasticity is the ability of cilia to repair. Ciliogenesis and ciliary length defects can recover *in vitro* and *in vivo* (*61–65*). Introduction of the wild-type copy of the mutant gene may rescue ciliogenesis defects. In some cases, ciliary defects of mutants may improve with the animal’s age and without the need for external intervention (64,65). The current known mechanisms involved in ciliary plasticity include an IFT kinesin, heat shock factor HSF-1, proteostasis and the unfolded protein response pathways, and ciliary GTPases involved in lipidated protein trafficking in cilia (64,65). Ciliary plasticity mechanisms are clinically relevant since ciliopathies such as polycystic kidney disease and retinal degenerative disorders are progressive and may amenable to rescue by a wild-type copy or other mechanisms (54,63). However, our knowledge of ciliary plasticity mechanisms - including ultrastructural plasticity - remains poor. Since defects in the TZ are reversible and the TZ allows incorporation of proteins into pre-assembled TZs (56,66), closing this knowledge gap has enormous therapeutic implications.

IL2Q neurons present an excellent model to identify mechanisms controlling ciliary plasticity. In IL2Q cilia, plasticity occurs in the form of TZ reorganization and MT structural rearrangements. Importantly, this TZ reorganization is not coupled to ciliogenesis and ciliary elongation, which allows the identification of molecules specifically involved in ciliary ultrastructural plasticity. Our identification of RAB-28, a ciliary GTPase implicated in human ciliopathies, as a plasticity regulator, demonstrates the utility of IL2Q cilia to discover conserved ciliary plasticity mechanisms.

The IL2 neurons are also useful to study how ciliary plasticity is modulated in different developmental and environmental contexts. IL2 cilia undergo reversible length changes when the animal enters the dauer stage, a diapause that allows survival in harsh environments. IL2 cilia shorten during dauer, are covered by the cuticle, and do not protrude into the environment; IL2 cilia elongate upon reintroduction of dauer animals to food and return to larval development (31,67). The mechanisms that regulate this form of signal-induced developmental ciliary plasticity remain unknown. Signaling programs involved in dauer entry, maintenance, and exit may play a role in this process. In reproductively growing animals, IL2 cilia are directly exposed to the environment. Environmental conditions influence tubulin glutamylation levels in amphid cilia of *C. elegans* (68). We hypothesize that environmental and developmental conditions influence glutamylation state in IL2 cilia. Examining the relationship between IL2 ciliary length, dauer, glutamylation, and the environment will provide insight to signal-induced ciliary plasticity mechanisms.

Several questions remain and new questions arise. Is ciliary specialization an ongoing process that is not only established but continuously maintained? Our data indicates that TZ specialization requires continuous maintenance. How is IFT affected by the variability in the individual MTs of IL2Q cilia? What is the relationship between the ciliary membrane and TZ specialization and reorganization? Is TZ specialization connected to cellular function? Deviations from canonical 9+0 MT arrangements are evolutionarily conserved, including in mammalian primary cilia (21–23,25–28,69–71). Answers to these questions will reveal how non-canonical ciliary ultrastructures are generated, how non-canonical variants impact ciliary transport and function, and how structural defects and ciliary repair impact human ciliopathies and potential therapies.

## Methods

***C. elegans* strains and maintenance** All strains were cultured and maintained similarly to the procedures described in Akella, Carter et al.2020. The strain *him-5 (e1490)* was used as the wild-type control. All strains were maintained at 20°C. Reporter strains were crossed into mutant backgrounds and desired cross-progeny were identified by following visual markers and by standard PCR genotyping. Transgenic animals were obtained by injecting constructs containing fluorescent protein fusions of genes of interest into young adult hermaphrodites. Desired transgenic lines were identified based on the presence of a co-injection marker, PCR and sequencing-based confirmation of the presence of the fluorescent tag, and an absence of off-target mutations after homology directed repair..

### Strain list

CB1490 *him-5(e1490) V*

PT2102 *pha-1(e2123) III; him-5(e1490) V; myEx686 [klp-6p::GFP::gKLP-6_3’UTR + pBX]*

PT2106 *tba-6(cxP4018) I; myIs1 pkd-2(sy606) IV; him-5(e1490) V*

PT3189 *rab-28(tm2636)IV; him-5(e1490) V*

PT3874 *my149 [mSc::TBA-6]I; him-5(e1490) V*

PT2679 *him-5(e1490)V; myIs23 [cil-7p::gCIL-7::GFP_3’UTR+ccRFP] X*

PT2925 *tba-6(cxP4018) I; him-5(e1490) V; myIs23 [cil-7p::gCIL7::GFP_3’UTR+ccRFP] X*

PT3265 *rab-28(tm2636) IV; him-5(e1490) V; myIs23[cil-7p::gCIL7::GFP_3’UTR+ccRFP] X*

### Fluorescence microscopy

In all cases where adults were imaged, L4 animals of desired genotypes were isolated the previous day to provide virgin adults on the day of imaging. For live imaging experiments, animals were placed on 4% agarose pads and immobilized in 10mM levamisole. Confocal imaging was performed on a Zeiss LSM880 point scanning confocal microscope equipped with an Airyscan detector and controlled by the Zen Black software. Images were acquired using a 40x/1.2 NA, water immersion or a 63x/1.4NA oil immersion objective. Confocal Z-projections were generated using Fiji, Zen Black, and Zen Blue software. All image analysis was performed using Fiji software.

For localization analysis of mSc::TBA-6 in IL2 neurons, confocal Z-projections were first assembled into montages on Fiji. Montages were examined to provide general descriptions of marker localization. Plot profiles generated using Fiji were also used to make assessments regarding enrichment in different subcellular regions within ciliated neurons.

Statistical analysis: Raw data was sorted and arranged using Microsoft Excel. Statistical analyses were done using GraphPad Prism V5. P values were indicated on all plots. Standard symbols were used to depict P values (* for p<0.05, ** for p<0.005, and *** for p<0.0005).

### Electron microscopy

TEM on wild-type control CB1490 and PT2106 adults was performed according to the method described in Silva *et al.* 2017. TEM on PT3189 adults was performed according to the method described in Akella *et al.* 2020. The IL2 ciliary data for wild-type and *tba-6* mutants in this paper was mined from the same animals imaged for CEM ciliary studies in Silva *et al*. 2017. This paper’s IL2 ciliary data for *rab-28* mutants was mined from the same animals imaged for CEM ciliary studies in Akella *et al.* 2020. All genotypes were synchronized as one-day-old virgins by isolating L4 males from hermaphrodites and not from each other the day before the high-pressure freeze fixation. Males were subjected to high-pressure freeze fixation using a HPM10 high-pressure freezing machine (Bal-Tec, Switzerland). Samples were freeze substituted in 2% osmium tetroxide, 0 - 0.2% uranyl acetate, and 2% water in acetone as the primary fixative, using RMC freeze substitution device (Boeckeler Instruments, Tucson, AZ, USA).

Samples were infiltrated with Embed 812 resin for two to three days before embedding and curing the blocks. 70 nm-thick serial sections were collected on copper slot grids with carbon-coated formvar film. Sections were post-stained with 2-4% uranyl acetate in 70% methanol, followed by washing and incubating with aqueous lead citrate. TEM images were acquired on either a Philips CM10 at 80 kV or a JEOL JEM-1400 at 120 kV. Tilt series were acquired on a JEOL JEM-1400 TEM using SerialEM software. These data were processed using etomo; tomograms were generated using the R-weighted back projection method. Serial tomograms were stitched to generate a volume spanning around 1.7 microns and annotated using IMOD software to generate models.

We used the posterior transition zone where microtubule doublets attached to the Y-links are clearly visible to count doublet microtubule numbers.

Transition zone lengths were calculated by counting the number of serial TEM images that contained Y-linked microtubules. Y-links that span at least half the thickness of a section would not be visible clearly on TEM images. Thus, the empirical error for these calculations can be approximated to +70nm.

We traced microtubules in tomogram volumes to assess microtubule termination, disassembly, and configurations. A- and B-tubules were identified based on whether the microtubule being traced was complete with 13 protofilaments or incomplete with 10 protofilaments at the transition zone level. Configurations where the origins could not be traced, or in an extremely rare case where only the B-tubule extended more distally, were noted as “other”. Configurations, where the MTs seemed to be associated with structural intermediates whose origins could be traced (at least partially to a doublet MT), were labeled “discontinuous”.

### Molecular biology

The mSc::TBA-6 reporter was generated using CRISPR/Cas9 mediated editing of the endogenous *tba-6* locus as described in (72) using a crRNA 5’CGAACAAUGCCACAAUACAAGUUUUAGAGCUAUGCU3’ and a double stranded DNA donor template that contained the fluorescent protein tag and the endogenous coding sequence. Animals expressing endogenously tagged TBA-6 were identified using PCR (forward primer sequence 5’ACAAATTGTCATCGTTCTCG3’ and reverse primer sequence 5’GTATGAACTCTAACTTCCATCTAAT3’). CRISPR lines of interest were also confirmed using sequencing-based detection of the mScarlet tag that was introduced.

## Acknowledgements

This work was supported by National Institutes of Health (NIH) DK059418, DK116606 and NS120745 (M.M.B) and by R24 OD010943 (D.H.H). Electron microscopy conducted at the Einstein Analytical Imaging Facility was supported by P30CA013330 and SIG #1S10OD016214-01A1. We thank Gloria Androwski and Helen Ushakov for their excellent technical assistance. We thank members of the Barr lab and the Rutgers *C. elegans* community for their feedback and constructive criticism throughout the project. We thank Wormbase and WormAtlas for online resources. JSA is grateful to Anne Norris and Julie Van De Weghe for careful reading of the manuscript. We thank Leslie Gunther-Cummins and Xheni Nishku at AECOM for assistance with high pressure freeze fixation. We also thank the National BioResource Project (Tokyo Women’s Medical College, Tokyo, Japan) and *Caenorhabditis* Genetics Center (CGC) for strains. The CGC is supported by the National Institutes of Health - Office of Research Infrastructure Programs (P40 OD010440).

## Author contributions

Conceptualization: J. S. A. and M. M. B.; Methodology, investigation and analysis: J. S. A., M. 1. S. S, K. C. Q. N., D. H. H. and M. M. B.; Manuscript writing: J. S. A., M. S. S and M. M. B

## Figure legends

**Video 1 (Related to Figure 1). Electron tomography-based model of an IL2Q cilium in wild-type animals.** Movie generated from serial tomograms of an IL2Q cilium in wild-type animals. Color code is the same as for figures 1F and 1G. Discontinuous singlet is marked in grey. Movie follows the IL2Q cilium along its entire length. Scale bar is 200 nm.

**Video 2 (Related to Figure 5) Electron tomography-based model of an IL2Q cilium in *rab-28 (tm2636)* animals.** Movie generated from serial tomograms of an IL2Q cilium in *rab-28 (tm2636)* animals. Color code is the same as for figures 4F and 4G. Scale bar is 200 nm.

---

**Figure 3 figure supplement 1.**
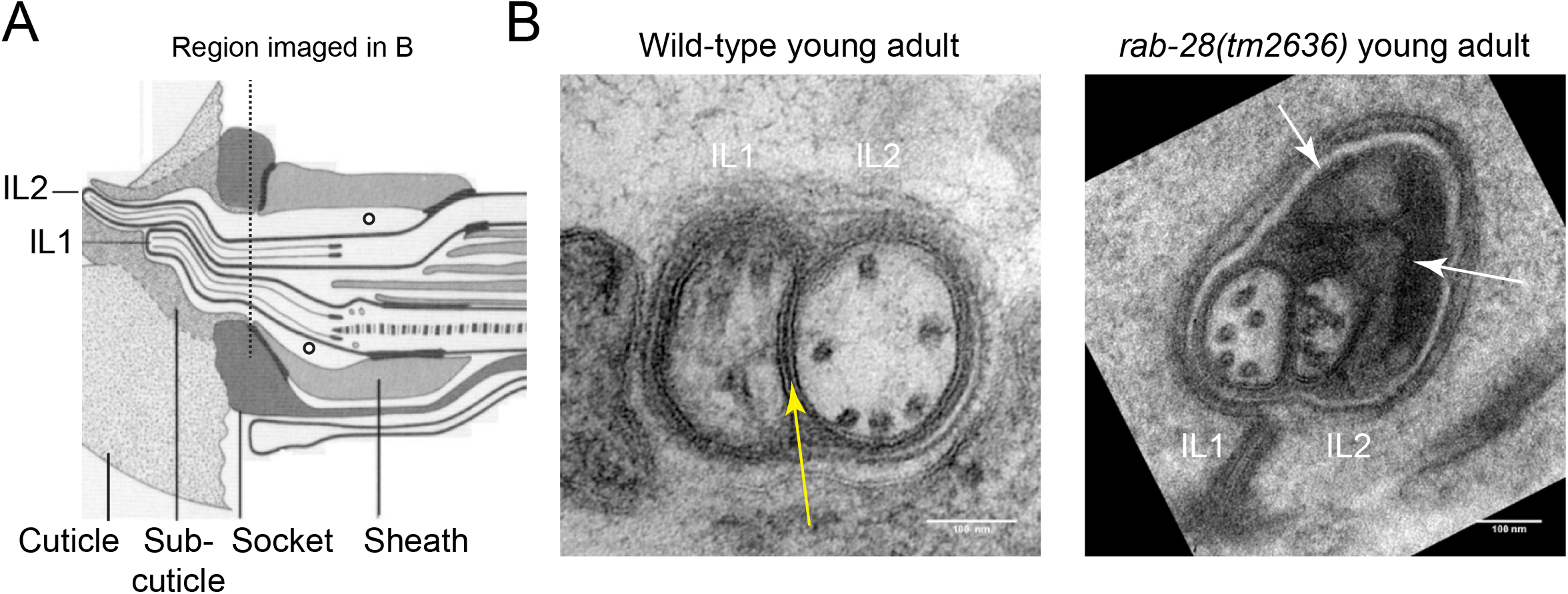
RAB-28 negatively regulates membrane shedding within inner labial sensory organs. (A) Same cartoon as in Figure 2A. Region marked with dotted lines represents the part of the sensory organ imaged by TEM in B. (B) Cross-section TEM of the inner labial sensory organ in the region distal to the TZ. Yellow arrow in wild-type panel points towards a gap junction between IL1 and IL2 cilia. *rab-28 (tm2636)* display an expansion of the inner labial sensory organ and accumulate an excess of membranous vesicles (white arrows). Scale bar is 100nm for both genotypes.

